## Supplemental Figures and Tables for "A novel attention mechanism for noise-adaptive and robust segmentation of microtubules in microscopy images"

### 1- Supplemental text

A- Generation of two synthetic datasets of fluorescence microtubules, called *MicSim_FluoMT*

To address the challenges of segmenting microtubules in noisy environments, we needed a dataset comprising hundreds of noisy images to train the neural network. However, generating such a real dataset is difficult, since collecting and annotating large numbers of images is costly and time-consuming, and high noise levels make accurate annotations unfeasible or imprecise. As a solution, we created two synthetic microscopy datasets of fluorescently labelled microtubules, along with corresponding binary masks (ground truth) using a two-step pipeline

*A1. Image dataset generation and annotation procedure*

We developed a two-step pipeline to generate synthetic images along with their corresponding ground-truths. The first step involved simulating the microtubule mitotic network in the *Caenorhabditis elegans* zygote, a well-established model for studying cell division [1]. For this, we used *Cytosim*, a widely used cytoskeletal simulation tool [2]. The simulations were parameterised using microtubule properties measured either *in vivo* or *in vitro*. We controlled key variables such as microtubule length, curvature, and density to reflect a range of biological scenarios (see table below. This approach allowed us to reproduce an astral microtubule network closely resembling that of the *C. elegans* embryo (Fig. S1A).

**Table:** Objects and their parameters of *Cytosim* simulations

| Object type | Characteristic parameters | Reference |
|---|---|---|
| Ellipse | Radii: 24.5 μm and 16.5 μm; Viscosity: draw from uniform distribution of values between 4 and 5 Pa.s. | Daniels *et al.*, 2006 |
| 2 solids (that mimic anterior and posterior centrosomes) | External force: Anterior: 180 pN; posterior: 500 pN. | Grill *et al.*, 2003 |
| 2 asters with two fibre types | Initial position: Anterior: -5.6 μm; Posterior 4.7 μm from ellipse centre. | *In vivo* lab measuremen ts |
| Fibre, type #1 (spindle) | Activity: classic; Number per aster: 20; Initial length 5.4 μm (for 10) and 6.8 μm (for 10); Rigidity: 50 pN.μm$^2$; Position: 60° fan distribution. | |
| Fibre, type #2 (astral) | Activity: dynamic; Number per aster: 65 to 85 (random choice); Initial length 8 ± 6 μm; Rigidity: draw from Gaussian distribution of mean equal 40 pN.μm$^2$ and variance equal 5 pN.μm$^2$; Position: 240° aleatory fan distribution; Growing force: 5 pN; Minimal length: 0.005 μm; Growing speed: 0.71 μm/s; Shrinking sped: -0.84 μm/s; Catastrophe rate: 0.05 (no force); 0.5 (stall force); Rescue rate: 0.15. | Dogterom *et al.*, 1997 ; Srayko *et al.*, 2005 |
| Single with hands (that mimic cortical force generators) | Activity: bind; Anchored to a fixed position; Unbinding rate: 0.1; Unbinding force: 5 pN. | 99.4% |

In the second step, we generated synthetic images from these simulations using *ConfocalGN*, an image generator that mimics confocal microscopy [3]. To closely match our real images, we adopted an empirical approach rather than modelling fluorescence intensity analytically. Specifically, we extracted fluorescence intensity distributions for both microtubule and background pixels from real deconvolved images of live dividing embryos acquired using an Airyscan microscope (Fig. S1B, D2, E2). Synthetic fluorescent images were then constructed in two stages: (1) by applying the Point-Spread-Function (PSF) blur and simulating photon noise, and (2) by adding background noise (Fig. S1C). This process resulted in what we refer to as the "*MicSim_FluoMT simple* dataset". To better replicate uneven fluorescence observed along astral microtubules, we created a second dataset in which fluorescence intensity gradually decreases toward the cell periphery. We refer to as the "*MicSim_FluoMT complex* dataset", which introduced greater segmentation difficulty, providing a more rigorous test for segmentation algorithms. The images closely resembled real microscopy data, accurately capturing microtubule fluorescence intensity (Fig. S1E1), background noise levels (Fig. S1D1) and filament morphology (Fig. S1C3). Finally, we created binary masks to precisely annotate microtubules in each image. These masks were automatically derived from the simulation data, ensuring perfect alignment between the synthetic microtubule images and their corresponding annotations. They served as ground truth for training supervised deep learning models and for evaluating segmentation performance.

*A2. Synthetic images: characteristics, visualisation and advantages*

We conducted 500 simulation runs and selected several time points from each to construct the two *MicSim_FluoMT* datasets, each consisting of 1192 synthetic 2D images. The diversity in fluorescence intensity, microtubule density, and background noise highlights the dataset's complexity and its usefulness for training and evaluating deep learning models for microtubule segmentation. Fig. S2 provides an overview of the *MicSim_FluoMT* datasets, showcasing representative examples of the synthetic images alongside their corresponding annotations. Both datasets are publicly available on Zenodo (DOI 10.5281/zenodo.14696279) [4].

Existing datasets of fluorescently labelled filaments are primarily based on real confocal microscopy images, which often lack annotations or include only partial annotations. In addition, the limited number of available images may not fully capture the morphological diversity present in biological samples. In contrast, our synthetic fluorescent microscopy datasets offer several distinct advantages that make them excellent resources for filament segmentation tasks in microscopy. (1) Controlled variability: Unlike real datasets, our synthetic data allows full control over key image parameters such as noise levels, fluorescence intensity, and filament morphology. This enables systematic evaluation of model performance under various controlled conditions. (2) Broad range of conditions: The dataset includes both simple cases (e.g., isolated microtubules in low-noise environment) and more complex scenarios (e.g., overlapping filaments with high noise levels), providing comprehensive test conditions for segmentation models. (3) Comprehensive annotations: Each image in the dataset is fully annotated at the pixel level – something rarely available in real datasets – allowing for precise training and robust evaluation of deep learning models. (4) Large dataset size: Our dataset contains a large number of images, ensuring that deep learning models can be trained effectively and tested across diverse conditions, helping to improve their generalization and robustness.

#### B- Generation of a real dataset of stained microtubules, called *MicReal_FluoMT*

To evaluate the performance of our new architecture in segmenting microtubules in real images, we created a dataset comprising images of microtubules stained with an anti-tubulin antibody (DM1A-AF488 conjugate) in *Caenorhabditis elegans* zygotes. To ensure variability in microtubule shapes and densities, which is important for effective model training, we included four experimental conditions: two targeting *zyg-8*$^{DCLK1}$ – a protein that binds to microtubules and regulates their rigidity – namely, *zyg-8(RNAi)* treated embryos and *zyg-8(or484ts)* heat-shocked mutants; and two control conditions, including RNAi control embryos and untreated heat-shocked embryos. We imaged fixed embryos using a confocal super-resolution microscope (Airyscan LSM980) and we acquired a stack of z-sections for each embryo. From each stack, we selected one or two z-sections that provided a clear visualisation of the astral microtubule network. When we selected two z-sections per image stack, we ensured they depicted distinct regions of the astral network. In total, we collected 49 images of *C. elegans* embryos, capturing two different stages of mitosis: metaphase and anaphase.

To annotate the microscopic images, we applied a three-step image processing pipeline, similar to the method described in [5]. (1) We performed an extended depth-of-field projection across 3 z-sections —specifically the section of interest along with the sections immediately above and below— to enhance microtubule continuity [6]. (2) We applied the Orientation Filter Transform to enhance filamentous pattern against noises [7]. (3) We applied the interactive machine-learning tool, *Ilastik,* to segment the astral microtubules [8]. During this semi-supervised segmentation, we manually annotated approximately 10 to 20 regions, labelling both microtubules and background in two to three embryos per condition. These annotations captured a range of intensities, contrasts, and microtubule morphologies, enabling the training of a segmentation model. This model was then applied to the remaining microscopy images to segment the astral microtubules in *C. elegans* embryos.

As a result, we obtained a dataset of 49 paired images, each consisting of a real microscopic image and a corresponding segmentation mask of the microtubules. We named this dataset "*MicReal_FluoMT*" and released it publicly on Zenodo (DOI 10.5281/zenodo.15852661) [9]. Of these, 19 images were used for training, 10 for validation, and 10 for testing. It is important to note that the semi-supervised segmentation occasionally identified structures outside the embryo that were not relevant to this study; however, these were retained in the segmented masks. Additionally, due to non-specific staining, the embryo periphery was often included in the segmentation masks. Some masks also contained small annotations within the embryo cytoplasm.

#### 2- Supplemental figures

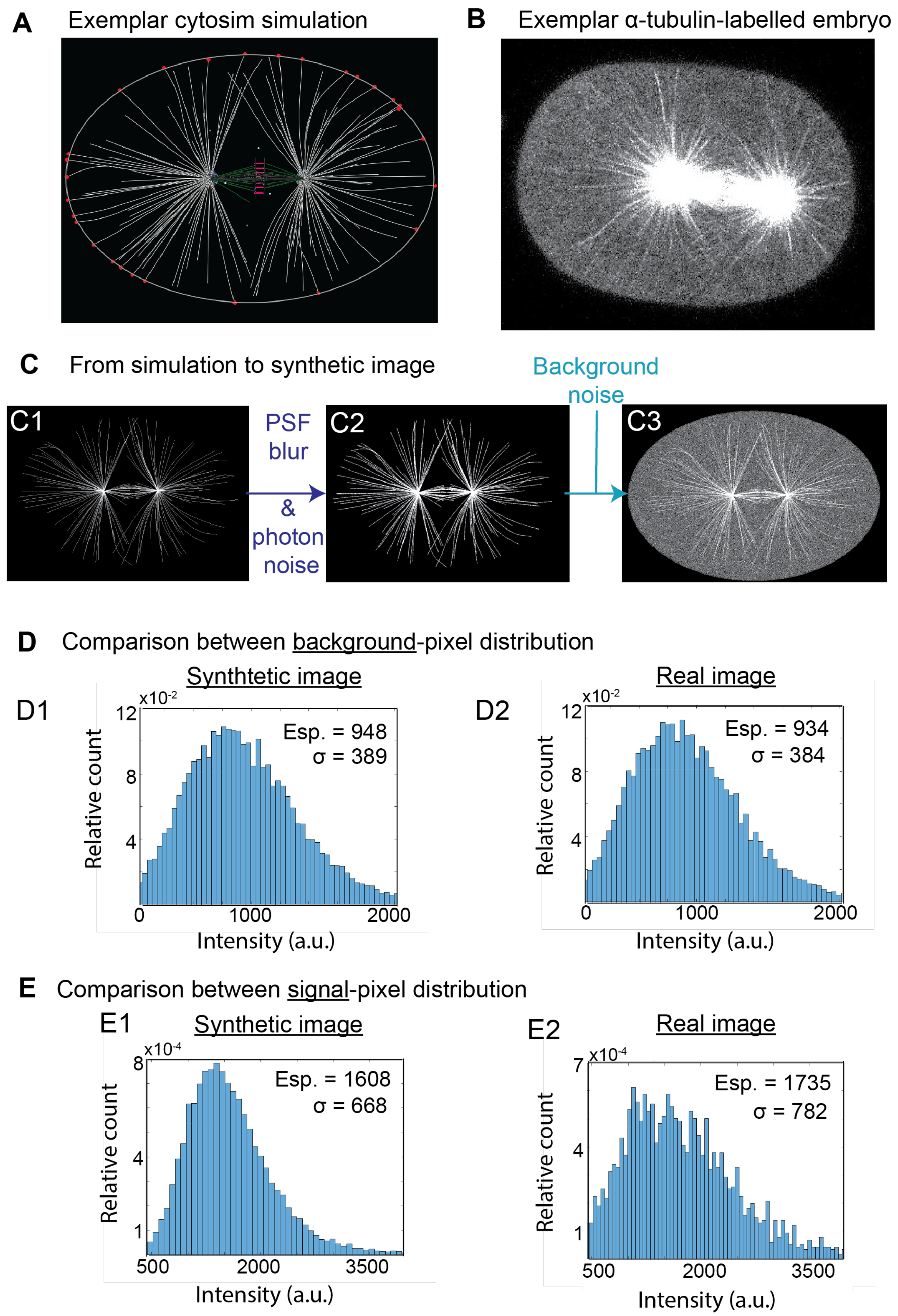

**Fig. S1: Generation of synthetic fluorescent images of microtubules for *MicSim_FluoMT* dataset, mimicking microtubule networks of *C. elegans* zygote visualised with fluorescently tagged tubulin.**

**(A)** Example of a *Cytosim* simulation showing astral microtubules (white lines); **(B)** Exemplar image of fluorescently labelled microtubules with YFP::$\alpha$-tubulin; **(C)** Two-stage generation of a synthetic image using *ConfocalGN*; **(D)** Comparison of background pixel intensity distributions between (D1) synthetic and (D2) real images; and **(E)** Comparison of microtubule pixel intensity distributions between (E1) synthetic and (E2) real images.

#### Representatitve images of MicSim FluoMT easy and complex datasets

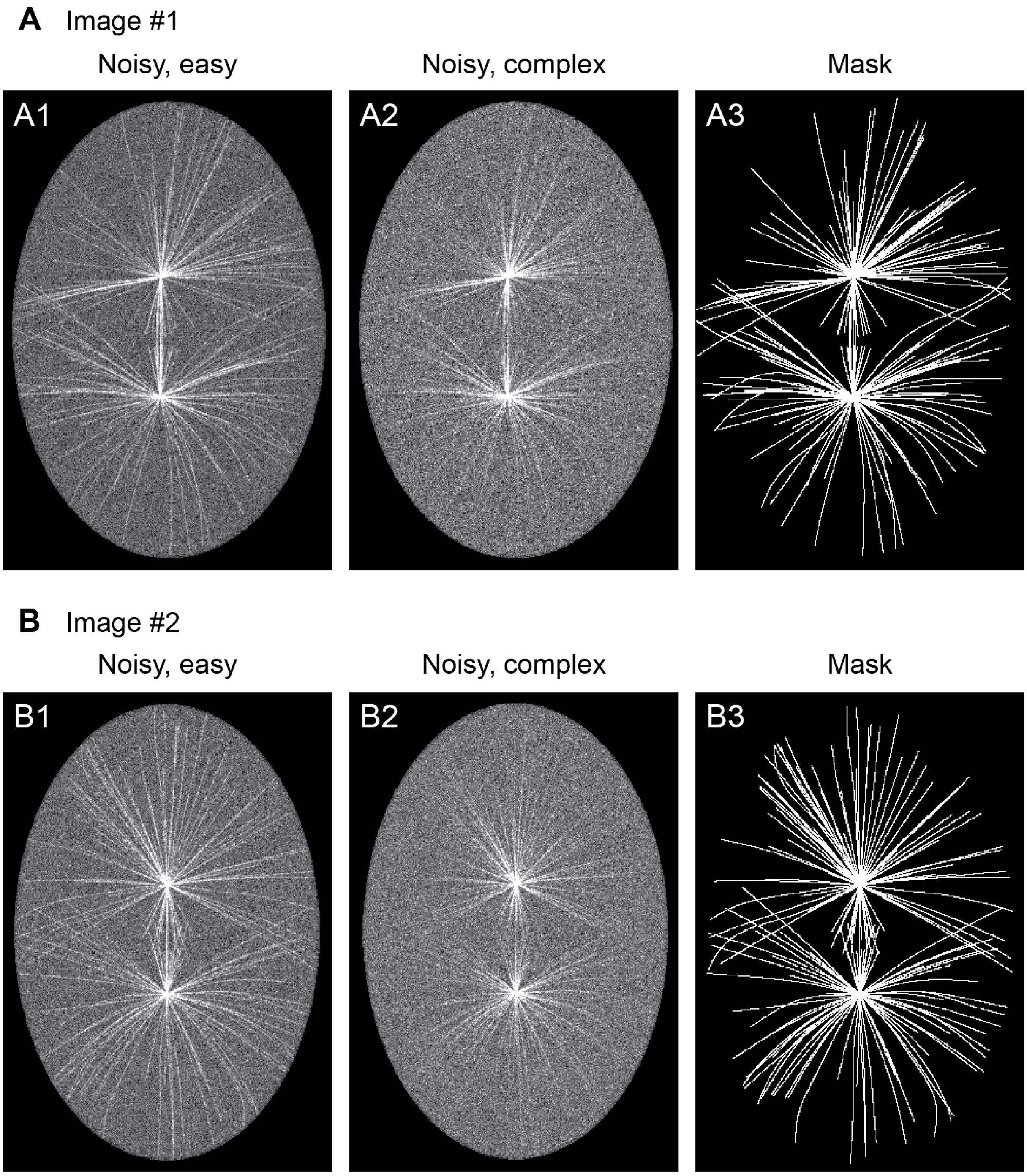

**Fig. S2: Overview of the two synthetic image datasets: *MicSim_FluoMT easy* and *MicSim_FluoMT complex*.**

Example images from *MicSim_FluoMT* datasets, with surrounding dark areas cropped to enhance filament visibility: (**A1, B1**) *Easy* images with uniform fluorescence along filaments and a signal-to-noise ratio (SNR) of (A1) 13.1 dB and (B1) 12.6 dB; the estimated noise levels *N*, calculated via 2D convolution (Eq. 2), are (A1) 0.336 and (B1) 0.346;(**A2, B2**) *Complex* images with uneven fluorescence and a SNR of (A2) 10.3 dB and (B2) 10.0 dB; the estimated noise levels *N* are (A2) 0.888 and (B2) 0.882; (**A3, C3**) Corresponding ground truth masks.

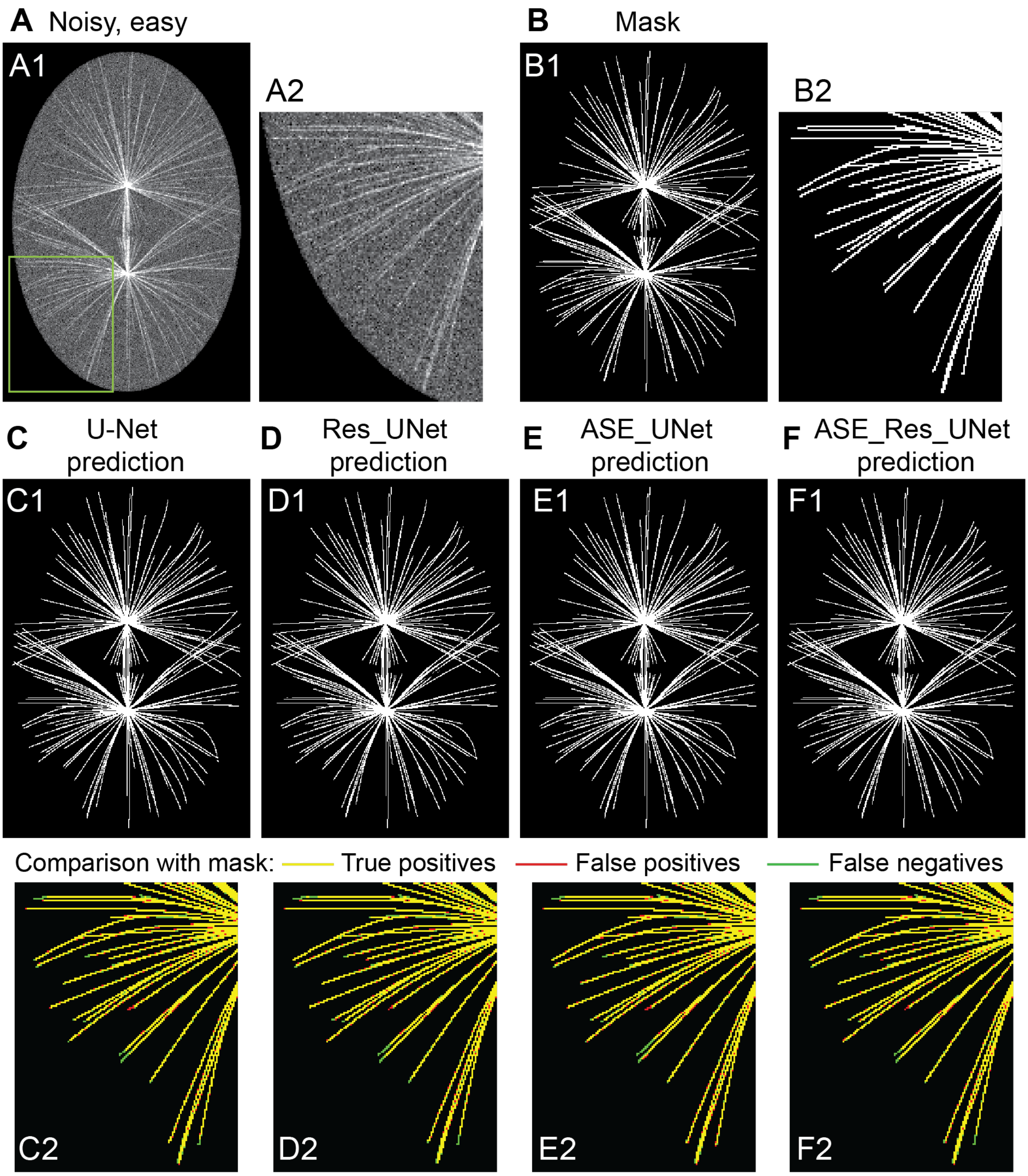

**Fig. S3: ASE_Res_UNet and its variants accurately segment microtubules in the *MicSim_FluoMT easy* dataset.**

Microtubule segmentation results obtained using ASE_Res_UNet and its variants on a sample test image from the *MicSim_FluoMT easy* dataset. (**A**) Input image; (**B**) Corresponding ground truth; (**C-F**) Predicted segmentations from (C) U-Net, (D) Res_UNet, (E) ASE_UNet, and (F) ASE_Res_UNet models. (A1-F1): whole simulated embryo; (A2-F2): zoomed-in regions of interest (ROI) to better highlight differences between model predictions and ground truth, with cropping window shown in green on panel A1; (C2-F2) composite ROI images showing true positives in yellow, false negatives in green, false positives in red, and true negatives in black.

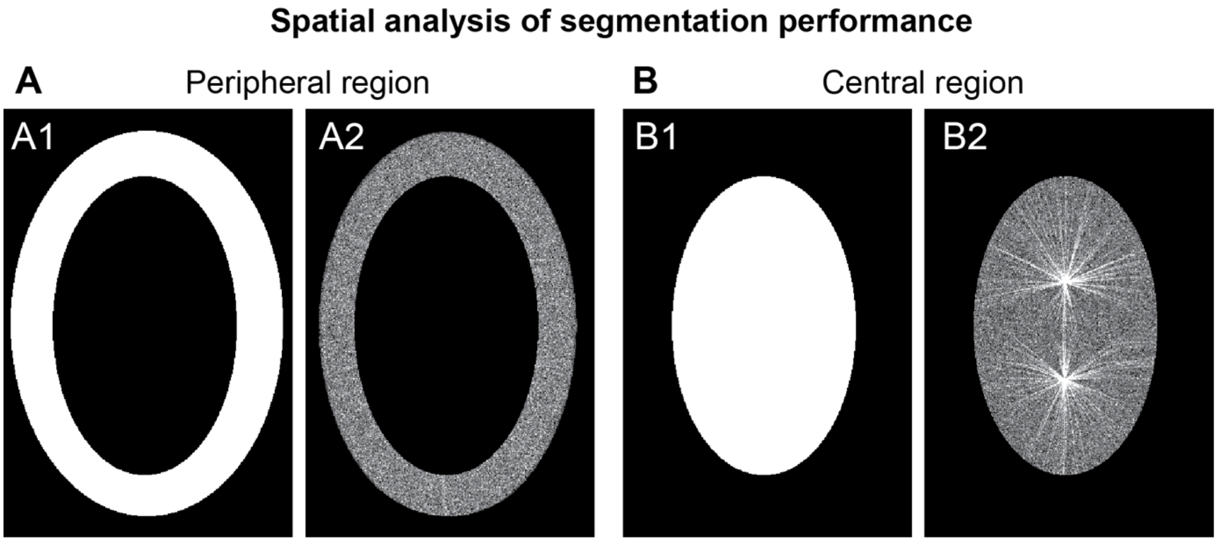

**Fig. S4: Definition of two embryo regions for spatial performance analysis in microtubule segmentation.**

Two regions of interest (ROI) were defined for spatial evaluation on the *MicSim_FluoMT complex* dataset: (**A**) peripheral region where microtubules have fluorescence intensities close to background noise; and (**B**) central region, where microtubules are more easily visible. (A1, B1) the masks of the ROI; and (A2, B2) ROIs overlaid on a representative test image.

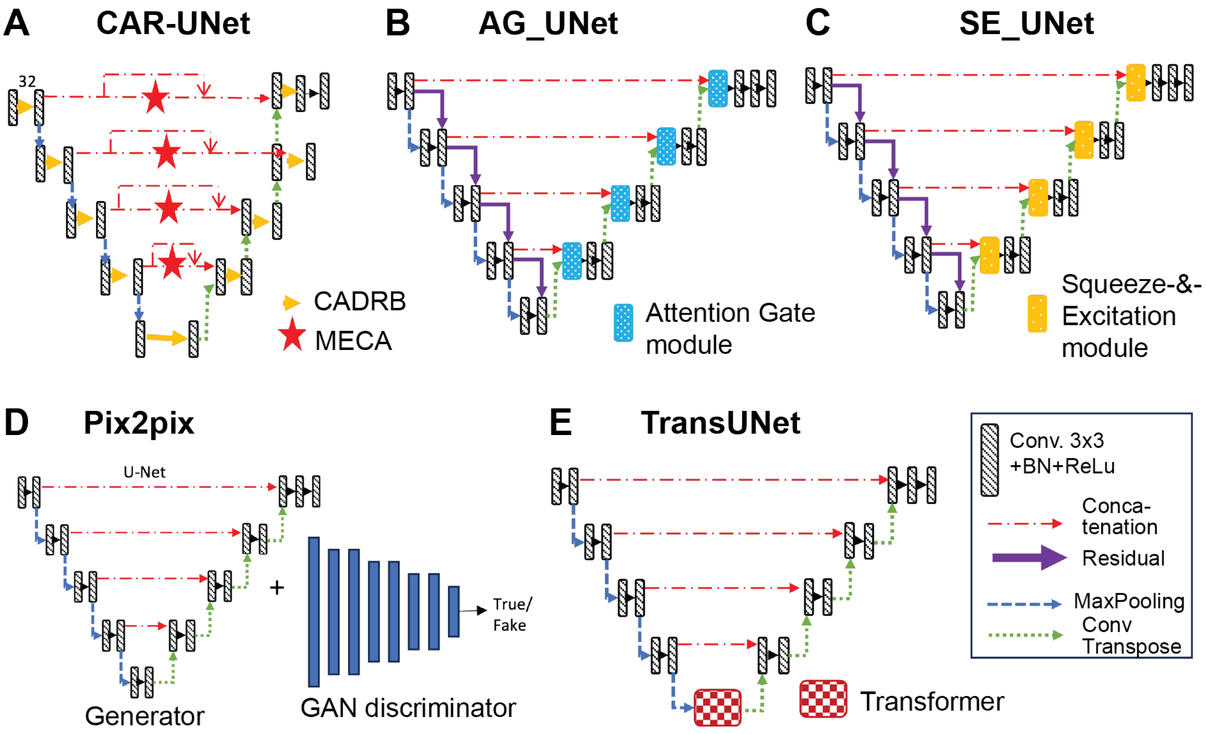

**Fig. S5: The five state-of-the-art architectures used for performance benchmarking in microtubule and vessel segmentation.**

Schematics of the architectures used to benchmark ASE_Res_UNet: (**A**) CAR-UNet, (**B**) AG_Res_UNet, (**C**) SE_Res_UNet, (**D**) Pix2pix, and (**E**) TransUNet.

### 3- Supplemental tables

| Metric | Description | Mathematical Formula | Relevance |
|---|---|---|---|
| **Dice** | Measures the overlap between predicted and ground truth masks. | $\frac{2TP}{2TP + FP + FN}$ | Good at evaluating segmentation where the positive class (e.g., filaments) is underrepresented; sensitive to boundary errors. |
| **IoU (Jaccard)** | Intersection over union between predicted and true regions. | $\frac{TP}{TP + FP + FN}$ | Stricter than Dice by penalizing more FP and FN; spatially accurate for verifying segmentation quality. |
| **Sensitivity** | Ability to correctly identify positive (foreground) pixels. | $\frac{TP}{TP + FN}$ | Crucial to avoid false negatives (i.e., missed curvilinear structures); ensures actual structures are captured. |
| **Precision** | Fraction of predicted positive pixels that are correct. | $\frac{TP}{TP + FP}$ | Important in noisy data; ensures that detected structures are truly relevant. |
| **MCC** | Correlation between predictions and true labels across all classes. | $\frac{TP \cdot TN - FP \cdot FN}{\sqrt{(TP + FP)(TP + FN)(TN + FP)(TN}}$ [illegible] | Strong measure under class imbalance; reflects overall prediction quality. |
| **PR AUC** | Area under the precision-recall curve. | $\int_0^1 \text{Precision(Recall)}\ d(\text{Recall})$ | More suitable than ROC AUC for imbalanced data; focuses on relevance of predicted positive structures. |

**Table S1**: **Six selected metrics for assessing segmentation performance in the presence of background class dominance**.

The table provides the description, formula and relevance of the following metrics: the Dice coefficient, Intersection Over Union (IoU), Sensitivity, Precision, Matthews Correlation Coefficient (MCC), and Area Under the Precision-Recall Curve (PR AUC). TP, FP, FN, and TN denote True Positives, False Positives, False Negatives, and True Negatives, respectively.

| Loss Function | Formula | Description | Relevance |
|---|---|---|---|
| **Binary Cross Entropy (BCE)** | $L_{\text{BCE}} = -\frac{1}{N}\sum_{i=1}^{N}[y_i \log(\hat{y}_i) + (1-y_i)\log(1-\hat{y}_i)]$ | Standard loss for binary classification, treats all pixels equally. | Works well on balanced datasets but struggles with background-foreground imbalance typical in filamentous structures. |
| **Weighted Cross Entropy (WCE)** | $L_{\text{WCE}} = -\frac{1}{N}\sum_{i=1}^{N}\left[\begin{matrix} w_1 \times y_i \log(\hat{y}_i) \\ + w_0 \times (1-y_i)\log(1-\hat{y}_i) \end{matrix}\right]$ | Extension of BCE, assigns higher weight to underrepresented class (e.g., filaments). | Helps mitigate foreground-background imbalance, shown effective in noisy microscopy settings. |
| **Focal Loss** | $L_{\text{Focal}} = -\alpha(1-\hat{y}_i)^{\gamma} y_i \log(\hat{y}_i)$<br>With α = 0.25 and γ = 2. | Focuses training on hard-to-classify examples by down-weighting easy ones. | Useful for datasets with extreme class imbalance or fine structures; however, tuning $\gamma$ and $\alpha$ is critical and does not promote structural accuracy. |
| **Dice Loss** | $L_{\text{Dice}} = 1 - \frac{2\sum y_i \hat{y}_i + \epsilon}{\sum y_i + \sum \hat{y}_i + \epsilon}$ | Directly optimizes for overlap between prediction and ground truth. | Helps preserve thin structures; however, may not capture fine topology and object boundary. |
| **Hausdorff Distance Loss** | Based on the Hausdorff distance:<br>$H(A,B) =$<br>$\max\{\sup_{a\in A}\inf_{b\in B} d(a,b), \sup_{b\in B}\inf_{a\in A} d(a$ | Penalizes segmentation errors based on spatial boundary discrepancies. | Strongly penalizes boundary mismatches, useful for assessing geometric accuracy of curvilinear shapes; however, limitations in case of class imbalance. |

**Table S2: Loss functions studied with their formula, description and relevance.**

$y_i$ is the ground truth label for the i-th pixel (0 for background, 1 for foreground) and $\hat{y}_1$ the predicted probability for the positive class (foreground) for the i-th pixel. N is the total number of pixels in the image. For WCE, $w_1$ and $w_0$ are the weights assigned to positive and negative classes, respectively. For the Focal loss, γ is a focusing parameter and α balances the importance of positive/negative examples. For Hausdorff Distance loss, *A* and *B* are sets of points on the contours of prediction and ground truth.

| Model | True Positive ↑ | True Negative ↑ | False Negative ↓ | False Positive ↓ |
|---|---|---|---|---|
| U-Net | 1 281 284 | 51 031 180 | 792 667 | 121 589 |
| Res_UNet | 1 306 883 | 50 998 731 | 767 068 | 154 038 |
| ASE_UNet | 1 455 872 | 50 902 052 | 618 079 | 250 717 |
| ASE_Res_UNet | 1 525 687 | 50 877 519 | 548 264 | 275 250 |

**Table S3: ASE_Res_UNet yields the highest number of True Positives and the lowest number of False Negatives, while maintaining a low number of False Positives.**

Confusion matrices of ASE_Res_UNet and its variant architectures, computed on the *MicSim_FluoMT complex* dataset (cumulative pixel count across 120 test images).

| Region | Model | Dice ↑ | IoU ↑ | Sensitivity ↑ | Precision ↑ | MCC ↑ | PR AUC ↑ |
|---|---|---|---|---|---|---|---|
| Central | U-Net | 0.8110 ± 0.0175 *** | 0.9797 ± 0.0016 *** | 0.7250 ± 0.0308 *** | **0.9216 ± 0.0142** *** | 0.8123 ± 0.0157 *** | 0.9256 ± 0.0095 * |
| | Res_UNet | 0.8117 ± 0.0164 *** | 0.9794 ± 0.0012 *** | 0.7365 ± 0.0269 *** | 0.9050 ± 0.0127 *** | 0.8112 ± 0.0153 *** | 0.9188 ± 0.0113 *** |
| | ASE_UNet | 0.8378 ± 0.0133 *** | 0.9812 ± 0.0011 *** | 0.8072 ± 0.0347 *** | 0.8736 ± 0.0344 ** | 0.8343 ± 0.0130 ** | 0.9271 ± 0.0106 |
| | ASE_Res_UNet | **0.8453 ± 0.0136** | **0.9817 ± 0.0009** | **0.8305 ± 0.0204** | 0.8608 ± 0.0098 | **0.8407 ± 0.0135** | **0.9297 ± 0.0101** |
| Peripheral | U-Net | 0.3595 ± 0.0526 *** | 0.9867 ± 0.0013 *** | 0.2315 ± 0.0427 *** | **0.8245 ± 0.0347 ***** | 0.4330 ± 0.0431 *** | 0.5605 ± 0.0534 ◊ |
| | Res_UNet | 0.3703 ± 0.0575 *** | 0.9866 ± 0.0012 *** | 0.2439 ± 0.0480 *** | 0.7882 ± 0.0360 *** | 0.4344 ± 0.0476 *** | 0.5391 ± 0.0556 *** |
| | ASE_UNet | 0.4448 ± 0.0541 *** | 0.9868 ± 0.0011 * | 0.3282 ± 0.0575 *** | 0.7115 ± 0.0659 * | 0.4778 ± 0.0454 *** | 0.5378 ± 0.0594 *** |
| | ASE_Res_UNet | **0.4997 ± 0.0488** | **0.9873 ± 0.0010** | **0.3933 ± 0.0520** | 0.6909 ± 0.0368 | **0.5175 ± 0.0433** | **0.5770 ± 0.0534** |

**Table S4: The performance gain of ASE_Res_UNet over its architectural variants is more pronounced in the peripheral region.**

Microtubule segmentation performances of ASE_Res_UNet and its variants on the *MicSim_FluoMT complex* dataset, evaluated in the peripheral and central regions of the embryo using various metrics (mean ± standard deviation over 120 test images). Bold values indicate the best performances for each metric. Statistical differences between ASE_Res_UNet and its variants are indicated only when significant (◊: $0.01 < p \leq 0.05$; *: $0.001 < p \leq 0.01$; **: $0.0001 < p \leq 0.001$; ***: $p \leq 0.0001$).

| Model | True Positive ↑ | True negative ↑ | False negative ↓ | False positive ↓ |
|---|---|---|---|---|
| CAR-UNet | 1 398 010 | 50 952 673 | 675 941 | 200 096 |
| AG_Res_UNet | 1 404 812 | 50 954 420 | 669 139 | 198 349 |
| SE_Res_UNet | 1 418 884 | 50 944 778 | 655 067 | 207 991 |
| TransUNet | 1 499 852 | 50 848 011 | 574 099 | 304 758 |
| Pix2pix | 1 412 662 | 50 543 648 | 661 289 | 609 121 |
| ASE_Res_UNet | 1 526 218 | 50 857 777 | 547 733 | 294 992 |

**Table S5: Compared to state-of-the-art models, ASE_Res_UNet yields the highest number of True Positives and the lowest number of False Negatives, while maintaining a low number of False Positives.**

Confusion matrices of ASE_Res_UNet and other advanced architectures, computed on the *MicSim_FluoMT complex* dataset (cumulative pixel count across 120 test images).

| Region | Model | Dice ↑ | IoU ↑ | Sensitivity ↑ | Precision ↑ | MCC ↑ | PR AUC ↑ |
|---|---|---|---|---|---|---|---|
| Central | CAR-UNet | 0.8309 ± 0.0137 *** | 0.9809 ± 0.0010 *** | 0.7802 ± 0.0207 *** | 0.8890 ± 0.0135 *** | 0.8279 ± 0.0134 *** | 0.9242 ± 0.0109 *** |
| | AG_Res_UNet | 0.8318 ± 0.0147 *** | 0.9810 ± 0.0010 *** | 0.7799 ± 0.0201 *** | **0.8913 ± 0.0137** *** | 0.8269 ± 0.0145 *** | 0.9265 ± 0.0115 ◊ |
| | SE_Res_UNet | 0.8338 ± 0.0160 *** | 0.9812 ± 0.0010 *** | 0.7858 ± 0.0241 *** | 0.8885 ± 0.0110 *** | 0.8307 ± 0.0159 *** | 0.9263 ± 0.0116 ◊ |
| | Pix2pix | 0.7549 ± 0.0166 *** | 0.9700 ± 0.0011 *** | 0.7712 ± 0.0141 *** | 0.7396 ± 0.0252 *** | 0.7473 ± 0.0166 *** | 0.8193 ± 0.0196 *** |
| | TransUNet | 0.8368 ± 0.0151 *** | 0.9808 ± 0.0009 *** | 0.8169 ± 0.0240 *** | 0.8579 ± 0.0084 ◊ | 0.8321 ± 0.0149 *** | 0.9221 ± 0.0118 *** |
| | ASE_Res_UNet | **0.8453 ± 0.0136** | **0.9817 ± 0.0009** | **0.8305 ± 0.0204** | 0.8608 ± 0.0098 | **0.8407 ± 0.0135** | **0.9297 ± 0.0101** |
| Peripheral | CAR-UNet | 0.4168 ± 0.0503 *** | 0.9869 ± 0.0011 * | 0.2905 ± 0.0460 *** | 0.7497 ± 0.0386 *** | 0.4628 ± 0.0423 *** | 0.5471 ± 0.0525 *** |
| | AG_Res_UNet | 0.4336 ± 0.0480 *** | 0.9870 ± 0.0011 | 0.3068 ± 0.0456 *** | **0.7538 ± 0.0565 *** ** | 0.4768 ± 0.0414 *** | 0.5663 ± 0.0551 |
| | SE_Res_UNet | 0.4386 ± 0.0540 *** | 0.9870 ± 0.0011 | 0.3136 ± 0.0519 *** | 0.7409 ± 0.0343 *** | 0.4781 ± 0.0457 *** | 0.5530 ± 0.0568 ** |
| | Pix2pix | 0.4092 ± 0.0390 *** | 0.9833 ± 0.0012 *** | 0.3580 ± 0.0418 *** | 0.4807 ± 0.0464 *** | 0.4101 ± 0.0384 *** | 0.4069 ± 0.0496 *** |
| | TransUNet | 0.4824 ± 0.0536 * | 0.9868 ± 0.0011 ** | 0.3816 ± 0.0589 | 0.6629 ± 0.0325 *** | 0.4989 ± 0.0472 * | 0.5450 ± 0.0574 *** |
| | ASE_Res_UNet | **0.4997 ± 0.0488** | **0.9873 ± 0.0010** | **0.3933 ± 0.0520** | 0.6909 ± 0.0368 | **0.5175 ± 0.0433** | **0.5770 ± 0.0534** |

**Table S6: ASE_Res_UNet outperforms state-of-the-art models in both central and peripheral regions, with the largest performance margin observed in the periphery.**

Microtubule segmentation performances of ASE_Res_UNet and other advanced architectures on the *MicSim_FluoMT complex* dataset, evaluated separately in embryo peripheral and central regions using various metrics (mean ± standard deviation over 120 test images). Bold values indicate the best performances for each metric. Statistical differences between ASE_Res_UNet and other architectures are indicated only when significant (◊: $0.01 < p \leq 0.05$; *: $0.001 < p \leq 0.01$; **: $0.0001 < p \leq 0.001$; ***: $p \leq 0.0001$).

#### 4- References

1. Hattersley N, Lara-Gonzalez P, Cheerambathur D, Gomez-Cavazos JS, Kim T, Prevo B, et al. Chapter 9 - Employing the one-cell C. elegans embryo to study cell division processes. In: Maiato H, Schuh M, editors. Methods in Cell Biology. 144: Academic Press; 2018. p. 185-231.
2. Nedelec F, Foethke D. Collective Langevin dynamics of flexible cytoskeletal fibers. New Journal of Physics. 2007;9(11):427.
3. Dmitrieff S, Nédélec F. ConfocalGN: A minimalistic confocal image generator. SoftwareX. 2017;6:243-7. doi: https://doi.org/10.1016/j.softx.2017.09.002.
4. Bouvrais H, Crespo M. MicSim_FluoMT: Two synthetic datasets of images of fluorescent microtubules (Ait Laydi et al., 2025). Zenodo; 2025.
5. Cueff L, Huet E, Schmitt L, Pastezeur S, Coquil M, Savary T, et al. Microtubule stiffening by doublecortin-domain protein ZYG-8 contributes to spindle orientation during <em>C. elegans</em> zygote division. bioRxiv. 2025:2024.11.29.624795. doi: 10.1101/2024.11.29.624795.
6. Forster B, Van De Ville D, Berent J, Sage D, Unser M, editors. Extended depth-of-focus for multi-channel microscopy images: a complex wavelet approach. Biomedical Imaging: Nano to Macro, 2004 IEEE International Symposium on; 2004: IEEE.
7. Sandberg K, Brega M. Segmentation of thin structures in electron micrographs using orientation fields. Journal of Structural Biology. 2007;157(2):403-15. doi: https://doi.org/10.1016/j.jsb.2006.09.007.
8. Berg S, Kutra D, Kroeger T, Straehle CN, Kausler BX, Haubold C, et al. ilastik: interactive machine learning for (bio)image analysis. Nature methods. 2019;16(12):1226-32. doi: 10.1038/s41592-019-0582-9.
9. Cueff L, Pecreaux J, Bouvrais H. MicReal_FluoMT: A dataset of microscopy images with stained microtubules (Ait Laydi et al., 2025). Zenodo; 2025.
